# Transport Pathways and Kinetics of Cerebrospinal Fluid Tracers in Mouse Brain Observed by Dynamic Contrast-Enhanced MRI

**DOI:** 10.1101/2022.11.03.515111

**Authors:** Yuran Zhu, Guanhua Wang, Chaitanya Kolluru, Yuning Gu, Huiyun Gao, Jing Zhang, Yunmei Wang, David L. Wilson, Xiaofeng Zhu, Chris A. Flask, Xin Yu

**Author notes:** Address correspondence to: Xin Yu, Sc.D., Wickenden 430, 10900 Euclid Avenue, Cleveland, OH 44106, USA.

## Abstract

Recent studies have suggested the glymphatic system as a key mechanism of waste removal in the brain. Dynamic contrast-enhanced MRI (DCE-MRI) using intracisternally administered contrast agents is a promising tool for assessing glymphatic function in the whole brain. In this study, we evaluated the transport kinetics and distribution of three MRI contrast agents with vastly different molecular sizes in mice. Our results demonstrate that oxygen-17 enriched water (H_2_^17^O), which has direct access to parenchymal tissues via aquaporin-4 water channels, exhibited significantly faster and more extensive transport compared to the two gadolinium-based contrast agents (Gd-DTPA and GadoSpin). Time-lagged correlation and clustering analyses also revealed different transport pathways for Gd-DTPA and H_2_^17^O. Furthermore, there were significant differences in transport kinetics of the three contrast agents to the lateral ventricles, reflecting the differences in forces that drive solute transport in the brain. These findings suggest the size-dependent transport pathways and kinetics of intracisternal contrast agents and the potential of DCE-MRI for assessing multiple aspects of solute transport in the glymphatic system.

## Introduction

Recent evidence suggests that the exchange of the cerebrospinal fluid (CSF) with the parenchymal interstitial fluid (ISF) occurs via a highly regulated, brain-wide pathway^1^. The glymphatic model proposes that CSF in the subarachnoid space is driven by cerebral arterial pulsation along the perivascular space surrounding penetrating arteries^2^, and its influx into the parenchyma is facilitated by the astroglial water channel aquaporin-4 (AQP4) located on the vascular endfeet^3^. The bulk flow from the influx of CSF into the parenchymal interstitium provides an efficient clearance route for metabolic by-products and other toxic wastes from the brain^4^. The transport of various CSF tracers has been studied extensively in rodent models to evaluate many pathophysiological factors that may impact CSF transport and CSF-ISF exchange, including the sleep-wake cycle, anesthesia, body postures, and cardiac function^5–10^. Further, impaired glymphatic function has been indicated in various disease conditions such as stroke, diabetes, traumatic brain injury, Alzheimer’s disease, and other dementias^11–16^.

Glymphatic flow and its dependence on AQP4 were first characterized in vivo by two-photon microscopy using fluorescent tracers with different molecular weights (MWs)^1^. Subsequent studies on four different lines of AQP4-knockout mice further confirmed the critical role of AQP4 in solute transport in the glymphatic system^3^. Dynamic contrast-enhanced MRI (DCE-MRI) provides the opportunity to assess both the kinetics and spatial distribution of CSF tracers in the whole brain^17,18^. Iliff and colleagues were the first to use DCE-MRI to evaluate the transport of MRI contrast agents in the glymphatic system in rat brains^19^. By comparing the transport of two gadolinium-based contrast agents (GBCAs) with different molecular sizes (Gd-DTPA, MW=938 Da; GadoSpin, MW=200 kDa), they showed that GadoSpin transport was confined to the subarachnoid space and CSF conduits, while Gd-DTPA was able to participate in CSF-ISF exchange. However, a limitation of using Gd-DTPA as a CSF tracer is that Gd-DTPA has limited penetration to the parenchyma because of its large molecular size, which may lead to an underestimation of the CSF-ISF exchange. Indeed, a recent study by Alshuhri et al compared the transport of Gd-DTPA and oxygen-17 (^17^O) enriched water (H_2_^17^O, MW=19 Da) in rats and reported significantly faster transport kinetics of H_2_^17^O, which is exchanged into the parenchyma directly via AQP4^20^. These studies suggest that the assessment of glymphatic function by DCE-MRI is dependent on the molecular size of the contrast agent, and using H_2_^17^O as a CSF tracer provides a unique opportunity to directly evaluate CSF-ISF exchange via AQP4.

Following these foundational DCE-MRI studies in rats, interest in evaluating the glymphatic function in mice grows rapidly due to the availability of genetically manipulated mouse models^3,6,11,21–25^. Assessing mouse glymphatic function by DCE-MRI has been challenged by the small size of a mouse brain and the limited volume (<20 μL) of fluids that can be delivered intracisternally without significantly altering the intracranial pressure (ICP)^15,26^. Early studies on mice used the protocol of administering contrast agents on the bench to allow close monitoring of the infusion process and to visually ensure proper sealing of the infusion site. While subsequent MRI scanning enabled assessing tracer transport at a later stage, such an approach inevitably missed the initial phase of contrast agent transport. More recently, Stanton et al and Gomolka et al successfully performed mouse DCE-MRI studies with in-scanner delivery of GBCA^9,27^. By monitoring the dynamics of GBCA transport for an hour, their results show that contrast enhancement peaked within 20 min of infusion in most of the regions characterized, highlighting the importance of delineating the kinetics of contrast agent transport at the early phase of infusion in mouse studies.

To build upon these prior studies, the goal of the current study was to evaluate the size-dependent transport kinetics and distribution of MRI contrast agents in the mouse glymphatic system. We first established and validated an intracisternal infusion protocol in mice that allowed the measurements of the entire time course of contrast agent transport for 2 hours. DCE-MRI studies were performed to compare the transport kinetics of Gd-DTPA, GadoSpin, and H_2_^17^O. Atlas-registered image analysis confirmed significantly faster transport of H_2_^17^O compared to Gd-DTPA, as well as drastically different transport kinetics of the three contrast agents from cisterna magna to the lateral ventricles. Furthermore, clustering analysis comparing the kinetic profiles of contrast agent induced signal changes showed a remarkable difference between H_2_^17^O and Gd-DTPA transport, suggesting different transport pathways for these two contrast agents.

## Results

### Transport of contrast agents in the whole brain

Fig. 1 shows representative sagittal slices of signal changes over the 2-hour DCE-MRI scans for each contrast agent. The signal changes showed clear differences in the transport dynamics of the three contrast agents (Figs. 1a-c). The time maximum intensity projection maps (tMIPs) further demonstrated the differences in contrast agent distributions during the entire time course of DCE-MRI scans (Figs. 1d-f). Following the infusion of the contrast agents at cisterna magna, contrast enhancement in the cerebellum region proximal to the infusion site can be observed within 5 min of infusion (CM in Fig. 1e). Subsequently, the transport of all three contrast agents to the fourth ventricle (V4), as well as along the subarachnoid space on the ventral surface of the brain, can be appreciated. Gd-DTPA was also transported into the parenchyma towards the dorsal direction after 15 min (Figs. 1a and 1d). In contrast, the transport of GadoSpin was primarily confined in the ventricles and the subarachnoid space (Figs. 1b and 1e). Compared to the two GBCAs, the transport of H_2_^17^O was more extensive and its distribution at the dorsal side of the brain was more pronounced (Figs. 3c and 3f). Finally, the downstream flow to the spinal cord (SC) was detectable for all three contrast agents.

**Fig. 1.**
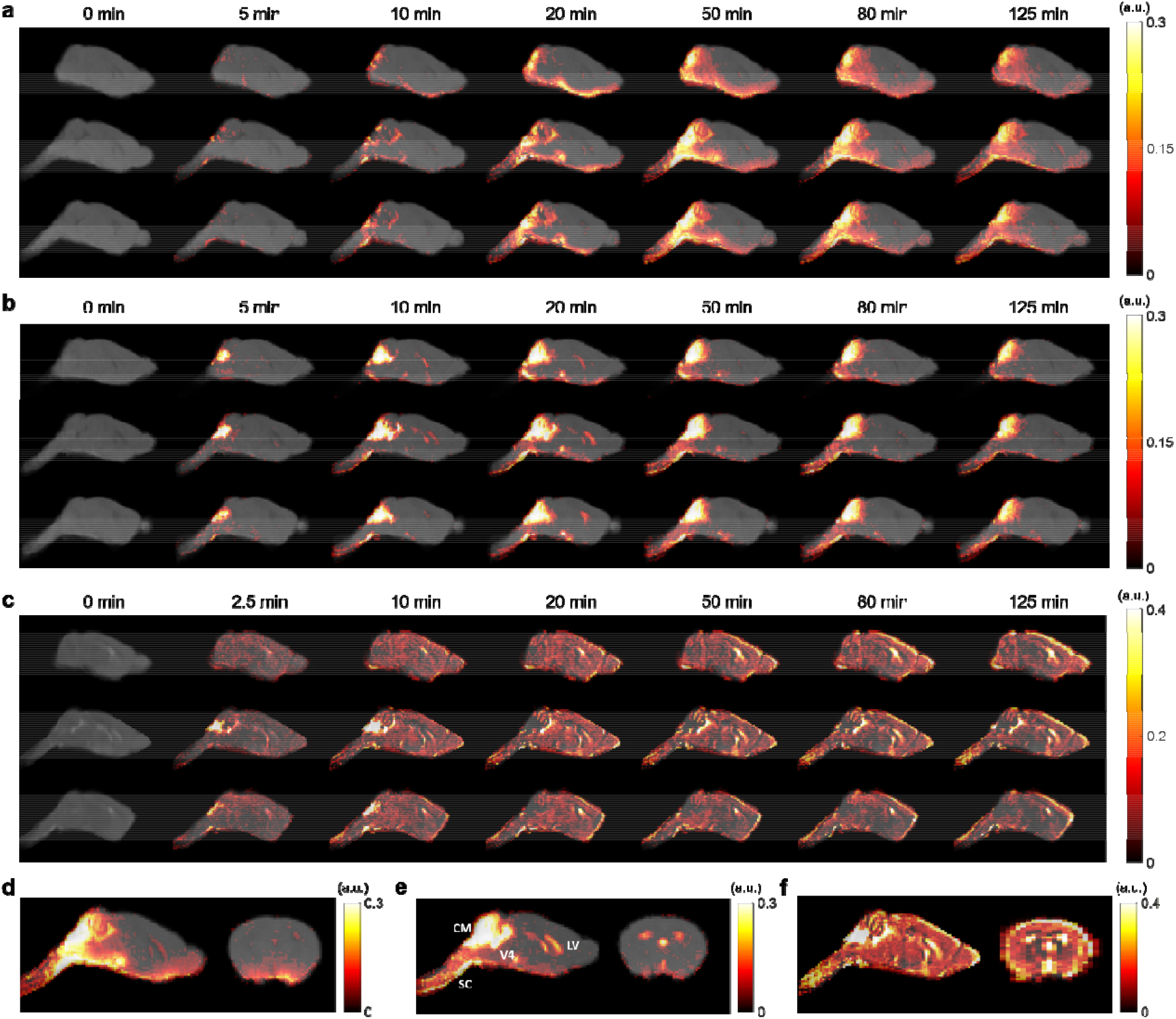
Contrast agent distribution. **a-c:** Representative sagittal views of group-averaged images, overlaid with signal changes from the baseline at selected time points, of mice infused with Gd-DTPA (**a**), GadoSpin (**b**), and H_2_^17^O (**c**), respectively. **d-f:** Time maximum intensity projection maps of representative sagittal and axial views of mice infused with Gd-DTPA (**d**), GadoSpin (**e**), and H_2_^17^O (**f**), respectively. CM, V4, LV, and SC in (**e**) indicate the contrast infusion site at cisterna magna, the fourth ventricle, the lateral ventricle, and the spinal cord, respectively.

### Dynamics of contrast agent transport

Shown in Figs. 2 to 4 and Fig. S1 are the time courses of signal changes in selected regions of interest (ROIs) covering the cerebellum and the ventral brain surface (Fig. 2), the deep brain (Fig. 3 and Fig. S1), the dorsal brain and the ventricular regions (Fig. 4). The dynamics of signal changes of the three contrast agents differ drastically in both the magnitude and the rate of change over time. In cerebellum (Fig. 2b), an ROI proximal to the site of contrast agent infusion, all three contrast agents showed rapid uptake, with H_2_^17^O being the fastest. While the accumulation of the two GBCAs remained high, H_2_^17^O exhibited very rapid clearance in the cerebellum, suggesting fast transport of H_2_^17^O from the infusion site. The transport of H_2_^17^O was also faster than that of Gd-DTPA in all the ROIs. In the deep brain and dorsal regions (Figs. 3 and 4), the transport of Gd-DTPA was further delayed compared to that in the ventral brain regions, with significantly reduced magnitude of signal enhancement that did not reach steady-state over a time period of ∼2 hrs. In contrast, H_2_^17^O transport in these regions was significantly faster and of a higher magnitude.

**Fig. 2.**
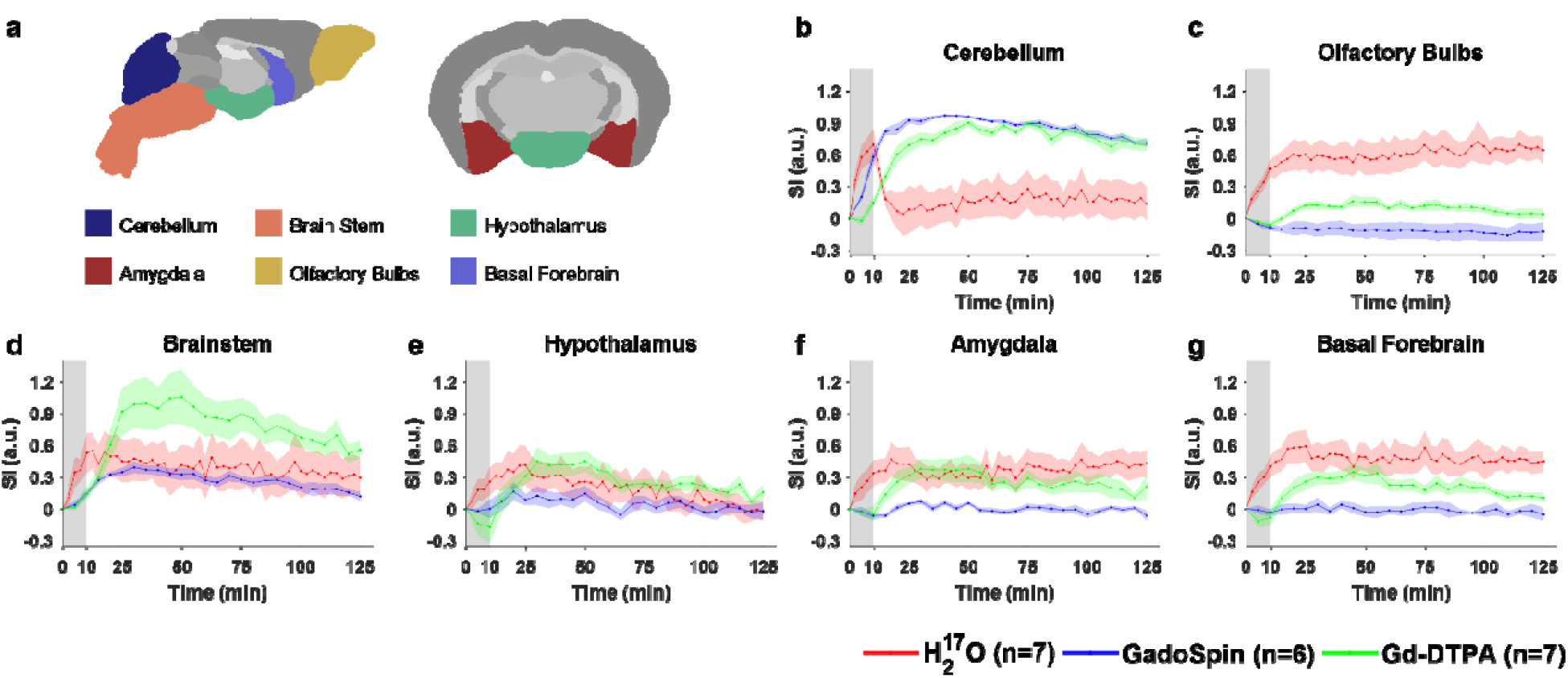
Contrast agent transport in cerebellum and ventral regions. **a:** Segmentation of selected ROIs. **b-g:** Time courses of signal changes in the selected ROIs. Gray bands indicate the time period of contrast agent infusion. Red, blue, and green lines represent the mean time courses of signal changes induced by H_2_^17^O, GadoSpin, and Gd-DTPA, respectively. Shaded areas represent standard errors.

**Fig. 3.**
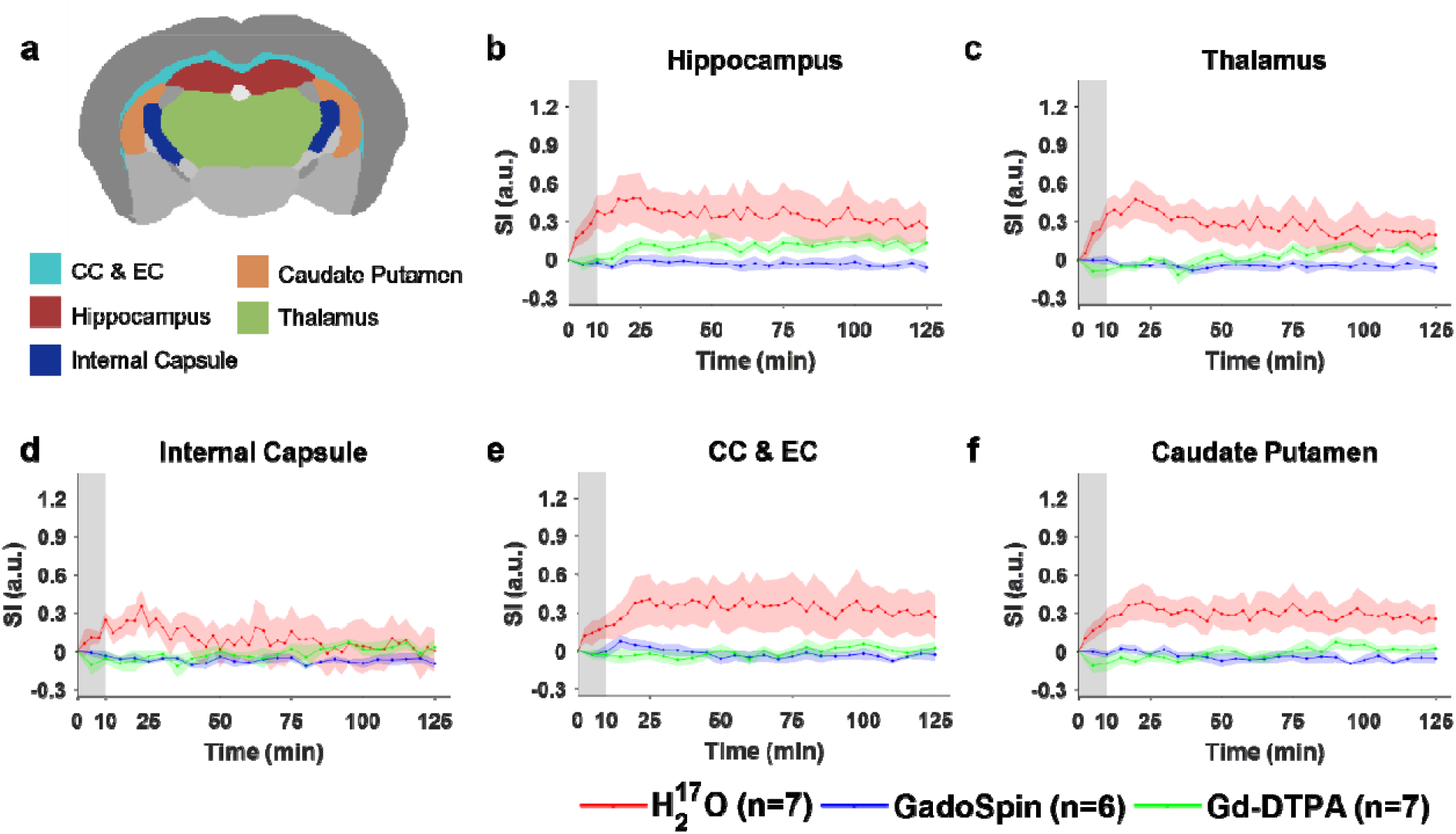
Contrast agent transport in deep brain regions. **a:** Segmentation of selected ROIs. **b-f:** Time courses of signal changes in the selected ROIs. Gray bands indicate the time period of contrast agent infusion. Red, blue, and green lines represent the mean time courses of signal changes induced by H_2_^17^O, GadoSpin, and Gd-DTPA, respectively. Shaded areas represent standard errors. CC & EC: Corpus callosum and external capsule.

**Fig. 4.**
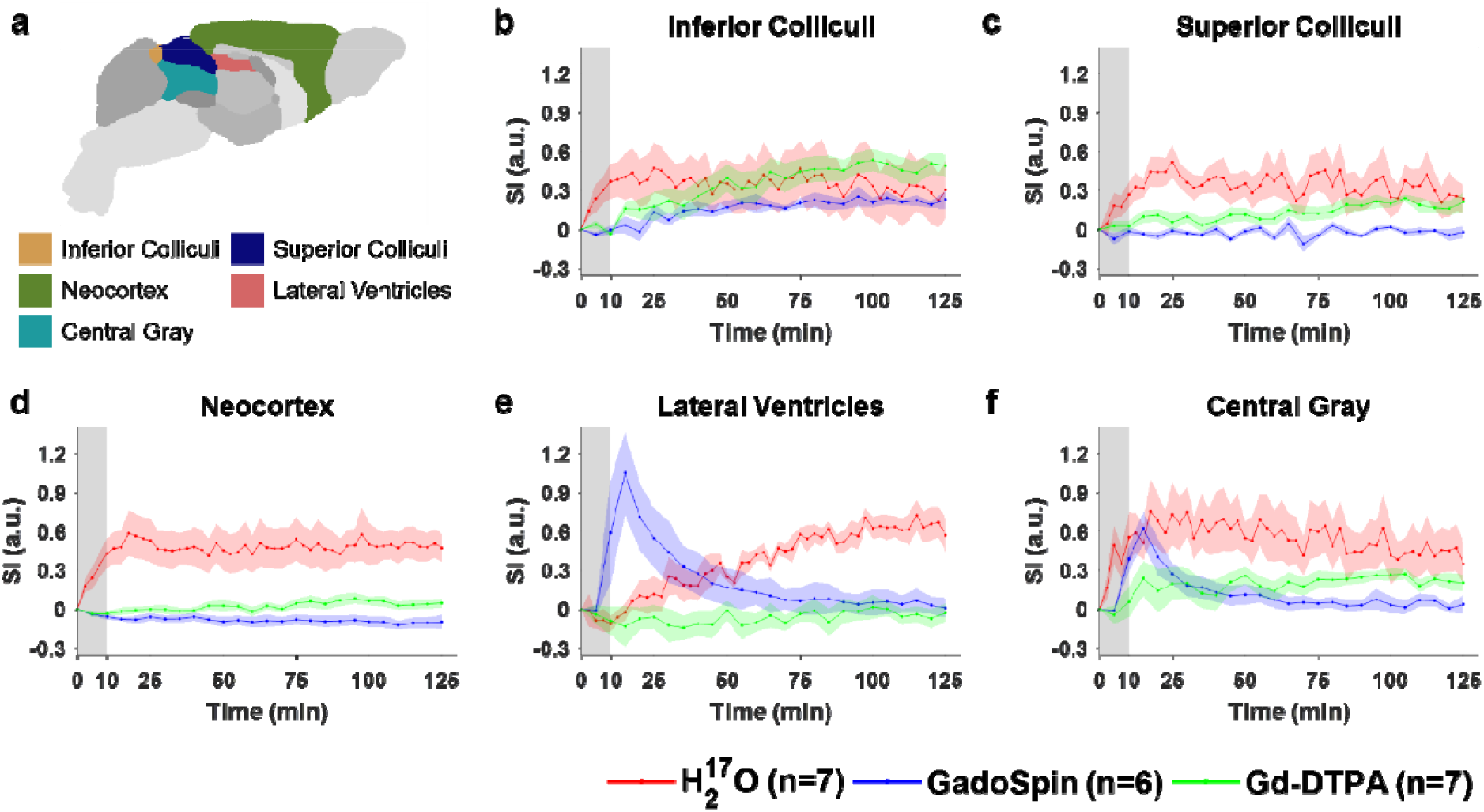
Contrast agent transport in dorsal brain, lateral ventricles and central gray regions. **a:** Segmentation of selected ROIs. **b-f:** Time courses of signal changes in the selected ROIs. Gray bands indicate the time period of contrast agent infusion. Red, blue, and green lines represent the mean time courses of signal changes induced by H_2_^17^O, GadoSpin, and Gd-DTPA, respectively. Shaded areas represent standard errors.

Compared to Gd-DTPA and H_2_^17^O, GadoSpin uptake only occurred in the ventricles and a few regions directly adjacent to the subarachnoid or large perivascular spaces, such as brainstem, inferior colliculi, and central gray. In the lateral ventricles (Fig. 4e), GadoSpin showed a prominent rapid uptake that peaked within 15 min after its infusion, followed immediately by a rapid clearance. Interestingly, a delayed but progressive accumulation of H_2_^17^O was observed, and the transport of Gd-DTPA to the lateral ventricles was negligible.

### Clustering of ROIs

Figs 5 and 6 show results of correlation-matrix-based hierarchical clustering analysis of Gd-DTPA and H_2_^17^O transport, respectively. The maximal cross-correlation coefficient (mCC) and the lag time corresponding to mCC between each pair of ROIs are shown in Figs. 5a and 6a. A large mCC value between two ROIs suggests a high similarity in the dynamic profiles of signal enhancement and the lag time indicates a relative delay in contrast agent transport between the two ROIs. Using 1-mCC as a measure of “dissimilarity” between two ROIs, hierarchical clustering analysis identified four clusters with a dissimilarity value of <0.4 were identified for both Gd-DTPA and H_2_^17^O. Bootstrap analysis with 1000 replications returned an approximately unbiased (AU) p-value of 0.92 to 1. These clusters are indicated by the rectangles in the correlation matrices in Figs. 5a and 6a, and their anatomical locations are outlined in Figs. 5b and 6b.

**Fig. 5.**
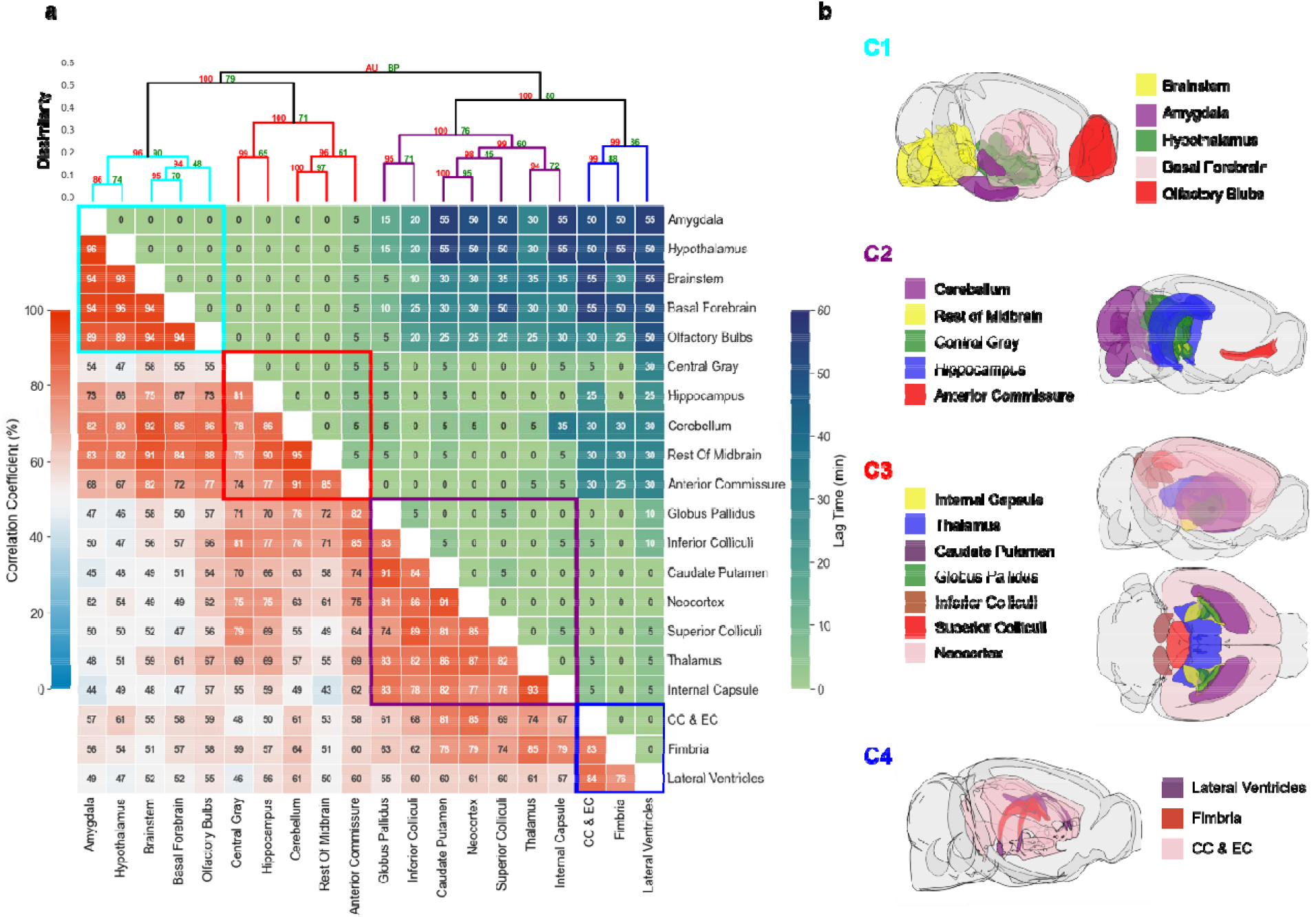
Correlation-matrix-based clustering analysis of Gd-DTPA transport. **a:** Matrix of maximal cross-correlation (lower left) and lag time (upper right), and cluster dendrogram with bootstrap analysis (top). Values at nodes are AU p-values (red, left) and BP values (green, right), respectively. Clusters with a dissimilarity value of <0.4 are indicated by the rectangles in (**a**). **b:** Outlines of ROIs in each cluster. CC & EC: Corpus callosum and external capsule.

**Fig. 6.**
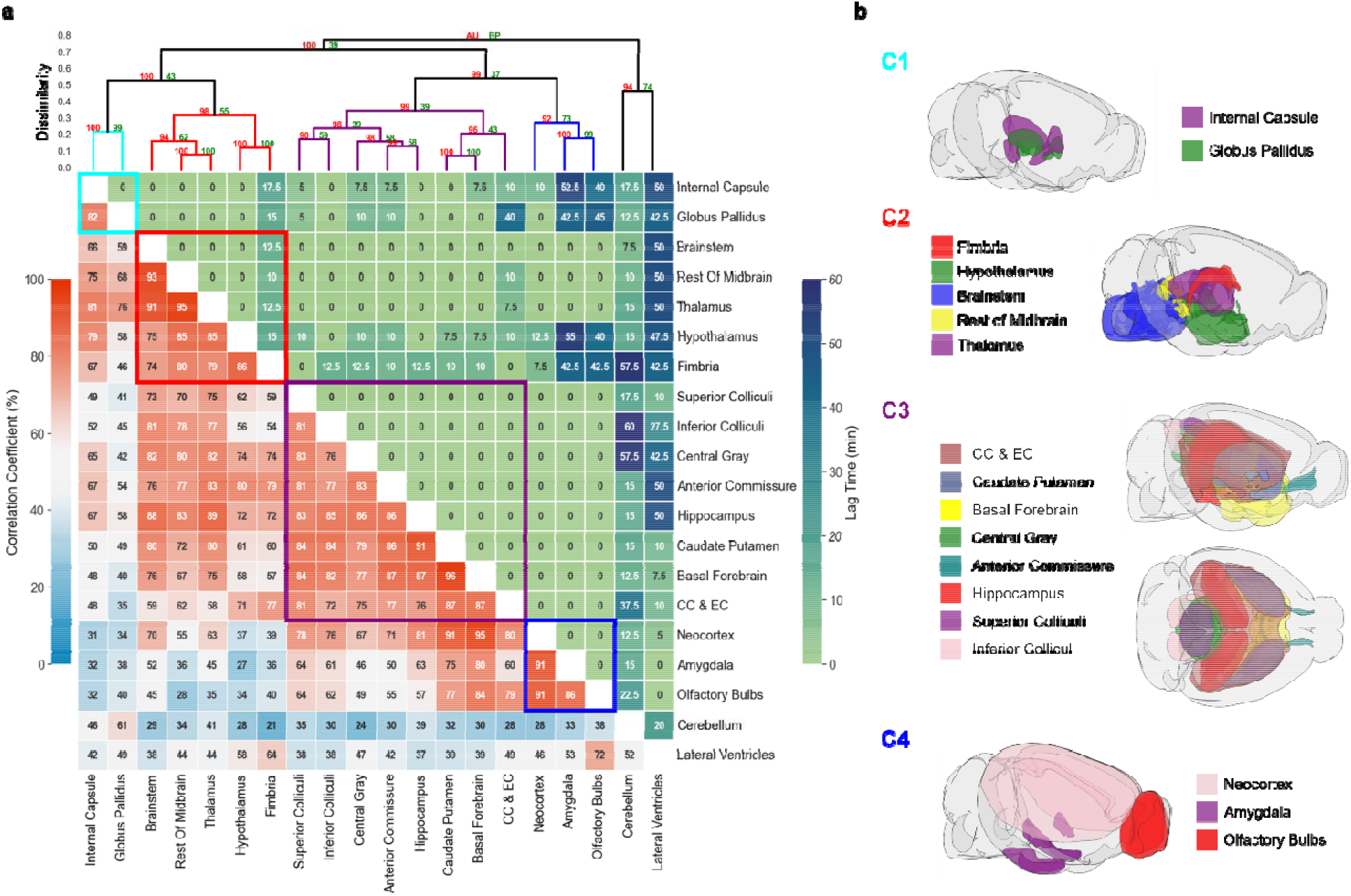
Correlation-matrix-based clustering analysis of H_2_^17^O transport. **a:** Matrix of maximal cross-correlation (lower left) and lag time (upper right), and cluster dendrogram with bootstrap analysis (top). Values at nodes are AU p-values (red, left) and BP values (green, right), respectively. Clusters with a dissimilarity value of <0.4 are indicated by the rectangles in (**a**). **b:** Outlines of ROIs in each cluster. CC & EC: Corpus callosum and external capsule.

For Gd-DTPA, all five ROIs along the ventral brain surface were classified into the same cluster (C1, cyan rectangle in Fig. 5a). The mCC values for this cluster were >93% for the four ROIs in the posterior to midbrain regions and decreased slightly to 89% in the olfactory bulbs in the anterior region. Further, the lag time was 0 among all these five ROIs, suggesting fast Gd-DTPA transport along the ventral surface of the brain. The remaining three clusters were grouped by their proximity to the subarachnoid space or ventral brain. Specifically, C2 (red) was anatomically adjacent to these regions, while C3 (purple) and C4 (blue) were located distally in the deep brain and dorsal regions. Correlation analysis of the ROIs in C3 and C4 with those in the ventral surface (C1) also showed the lowest mCC, as well as a longer lag time ranging from 25 to 55 min (Fig. 5a). In contrast, the lag time between ROIs in C1 and C2 was 0 except for anterior commissure (5 min).

H_2_^17^O transport showed distinctly different patterns in correlation matrix, lag time, and clustering from those of Gd-DTPA (Fig. 6a). The five ROIs along the ventral surface were classified into three different clusters (C2-C4) instead of one. The clusters were primarily grouped by their proximity to the infusion site, with two clusters (C1 and C2) located in the posterior to midbrain regions and the other two clusters (C3 and C4) located in the midbrain to anterior regions. Lag time between ROIs in the same cluster was 0 except for fimbria, the most anterior ROI in C2, which showed a lag time of 10-15 min from other ROIs in C2. Further, the lag time between two ROIs in different clusters was typically <15 min. Only a few ROIs showed a longer lag time compared to other ROIs, such as the cerebellum, lateral ventricles, amygdala, olfactory bulbs, corpus callosum and external capsule. The longer lag time between these ROIs can be attributed to the differences in the kinetic profiles rather than a delay in H_2_^17^O transport, as indicated by the very low mCC values (IZ40%) between these ROIs.

### Impact of H_2_^17^O exchange with blood

To evaluate the impact of the exchange of intracisternal H_2_^17^O with systemic circulation, blood samples were collected from mice at 20 min (n=3) and 125 min (n=4) after H_2_^17^O infusion, as well as from mice without H_2_^17^O infusion (n=3). T_2_ measurements of the plasma samples did not show significant differences among these three groups (Fig. S2), suggesting that H_2_^17^O exchange across the blood-brain barrier (BBB) did not significantly alter the T_2_ of circulating blood.

## Discussion

This study investigated the size-dependent transport of intracisternally administered CSF tracers in mouse brain using DCE-MRI. Consistent with previous observations in rats^19^, Gd-DTPA showed relatively fast transport kinetics along the ventral surface of the brain but delayed and reduced transport into the deep brain and dorsal regions. GadoSpin transport showed limited penetration to all parenchymal regions of the brain. Compared to the two GBCAs, the transport of H_2_^17^O was both faster and more extensive, which is also in agreement with a previous study in rats^20^. These findings were further supported by time-lagged cross-correlation analysis in which Gd-DTPA showed a much longer lag time between ROIs in the ventral surface versus those in the deep brain and dorsal regions as compared to H_2_^17^O. In addition, clustering analysis also showed different patterns of transport between Gd-DTPA and H_2_^17^O. Further, the three tracers also showed drastically different transport kinetics from cisterna magna to the lateral ventricles.

Previous studies of rat glymphatic function have employed pixel-based clustering analysis to identify regions with similar contrast enhancement features^19,28^. Using the area-under-the-curve in k-means clustering algorithm, Iliff et al demonstrated deeper penetration of Gd-DTPA to parenchymal tissues than GadoSpin^19^. In an effort to model the glymphatic system using local input function, Davoodi-Bojd et al used the derivatives of the time signal curve to identify clusters with similar profiles of contrast enhancement^28^. In the current study, we performed ROI-based hierarchical clustering analysis using mCC-derived parameters as a measure of “dissimilarity” between two ROIs or clusters. Such an approach allowed the identification of ROIs with similar kinetic profiles over the entire time course of imaging (2 hours). While lag time was not used as a parameter in clustering analysis, most of the ROIs within a cluster showed minimal lag time for both Gd-DTPA and H_2_^17^O, suggesting that this approach is robust in identifying ROIs along the transport pathway.

The clustering of Gd-DTPA transport was highly consistent with that by Iliff et al using a pixel-based clustering approach. In particular, all five ROIs along the ventral surface were identified as a single cluster while the remaining ROIs in the parenchymal tissues were clustered by their proximity to the subarachnoid space and ventral surface of the brain. More importantly, ROIs in these clusters also showed a prolonged lag time from the ROIs in the ventral cluster. These analyses suggest that Gd-DTPA transport occurred initially via the perivascular network, followed by its penetration into the parenchyma. In contrast, H_2_^17^O transport showed dramatically different clustering patterns compared to Gd-DTPA, with the five ventral ROIs distributed in three different clusters. In general, clustering of H_2_^17^O transport followed a pattern of posterior to anterior while Gd-DTPA was along the ventral surface towards the dorsal direction. These different clustering patterns reflect different transport pathways for H_2_^17^O because of its direct access to parenchymal tissues via AQP4.

A potential confounding factor in H_2_^17^O transport is the exchange of water across the BBB. Since systemic circulation is much faster than CSF flow^29,30^, its participation in H_2_^17^O transport may significantly accelerate the distribution of H_2_^17^O to the whole brain. To evaluate the impact of H_2_^17^O exchange across BBB, we measured the T_2_ of plasma collected at two time points, one at the time when H_2_^17^O-induced T_2_ change reached peak in parenchyma (20 min) and one at the end of the scan protocol (125 min). Our measurements showed no significant difference in plasma T_2_ at both time points compared to samples collected from mice without H_2_^17^O infusion. This lack of change in plasma T_2_ suggests that the intracisternal administration of a small volume of H_2_^17^O (10 μL) had a negligible effect on plasma H_2_^17^O concentration. Hence, it is unlikely that systemic circulation played a significant role in the observed fast H_2_^17^O transport in the current study. The participation of cerebrovasculature in H_2_^17^O transport needs more careful evaluation, as the possibility of H_2_^17^O entering the bloodstream near the infusion site and impacting signals in the cerebellum cannot be ruled out completely. However, considering that arteries in the cerebellum are not the feeding arteries to other regions of the brain and the very short transit time of cerebral blood in a mouse brain^31^, CSF-blood exchange would have to be extremely fast for cerebrovasculature to play a significant role in H_2_^17^O transport beyond cerebellum. Furthermore, Alshuhri et al reported that rats pretreated with an AQP4 inhibitor exhibited ∼80% reduction in H_2_^17^O transport into the parenchyma, providing strong evidence that AQP4 and the glymphatic system play a major role in H_2_^17^O transport to the parenchymal tissue^20^.

A recent study using confocal microscopy and fluorescein-labeled dextran (MW=500 kDa) as a CSF tracer has suggested that tracers administered via cisterna magna can be transported to the lateral ventricles via the interpeduncular cistern^32^. However, such observation was not unequivocally supported by a DCE-MRI study using Gd-DOTA (MW=661.8 Da) as the tracer, in which only one rat with an abnormally enlarged third ventricle exhibited Gd-DOTA uptake in the lateral ventricles^33^. In the current study, mice also showed negligible uptake of Gd-DTPA into the lateral ventricles. However, there was a rapid uptake of GadoSpin into the lateral ventricles that peaked at the end of its infusion, and it was immediately followed by a rapid clearance. Interestingly, despite its fast transport to the parenchymal tissues, H_2_^17^O showed a slow but progressive accumulation of H_2_^17^O in the lateral ventricles (Fig. 4e). Considering that cisterna magna is at the downstream of CSF circulation from the fourth ventricle^34^, direct transport of CSF tracers from cisterna magna to the lateral ventricles would require a reversal of the pressure gradient. The transport of GadoSpin into the lateral ventricles suggests that intracisternal infusion may have caused such a transient reversal of the pressure gradient between the cisterna magna and the lateral ventricles, and the rapid clearance of GadoSpin appears to be consistent with the restoration of the normal pressure gradient immediately after the infusion. On the other hand, the delayed transport of H_2_^17^O and the negligible transport of Gd-DTPA into the lateral ventricles suggest that the pressure gradient and the transport impedance for these two contrast agents may have favored their transport from the subarachnoid space into the parenchymal tissues. As such, the delayed H_2_^17^O uptake into the lateral ventricles may have occurred at the parenchymal-ventricular interface following its transport to the parenchyma. However, due to the slow and reduced penetration of Gd-DTPA into the parenchyma, Gd-DTPA did not reach the parenchymal-ventricular interface in a significant amount, leading to its negligible uptake into the lateral ventricles. Further investigations rooted in the fluid dynamics of CSF circulation are warranted to gain more quantitative insight of these observations.

There are several limitations in the current study. First, the use of T_1_- or T_2_-weighted images in DCE-MRI is a semi-quantitative approach such that signal changes from the baseline can only serve as a proxy of contrast agent concentration. While the concentrations of gadolinium ions (Gd^2+^) were matched for the two GBCAs, a direct comparison of contrast agent concentrations is not possible because of the nonlinear relationship between signal changes and contrast agent concentrations. As such, the current study focused on evaluating the kinetics of contrast agent induced signal changes. To account for inter-subject variations, we normalized the signal changes by using the maximal signal surrounding the infusion site as the common denominator. While this approach has enabled the comparison of relative signal changes in different ROIs, flux quantification is challenged by the nonlinear relationship between signal change and contrast concentration. Second, due to hardware limitations, a lower spatial resolution was used in the current study to achieve adequate temporal resolution, which has led to the pronounced partial volume effect in a few ROIs. For example, the large signal increase in the central gray from mice infused with GadoSpin (Fig. 4e) is likely caused by GadoSpin uptake in the fourth ventricle. Improving the spatial resolution will reduce the partial volume effect and enable the segmentation of smaller ROIs in future studies. Finally, previous studies have shown that general anesthesia with isoflurane alone can impair the circulation of CSF through the brain while low-dose isoflurane with supplementary dexmedetomidine or ketamine/xylazine can enhance glymphatic transport^5,6,9^. Hence, an anesthetic regime that enhances glymphatic function can be more desirable in studying solute transport in the brain, especially when using tracers with a large molecular size such as Gd-DTPA.

In summary, the current study established an experimental protocol that enabled DCE-MRI measurements of the entire time course of contrast agent transport in the mouse glymphatic system. Comparison of the transport kinetics and distribution of three MRI contrast agents with different molecular sizes showed drastically different transport profiles and clustering patterns, suggesting that the transport pathways and kinetics in the glymphatic system are size-dependent.

## Materials and Methods

### Animals

The animal protocol was approved by the Institutional Animal Care and Use Committee of Case Western Reserve University. The experiments were performed on 13- to 15-week-old male C57BL/6J mice (Jackson Laboratories, Bar Harbor, ME, US). The average body weight at the time of MRI scan was 29.4 g. The animals were housed in a temperature- and humidity-controlled environment with ad libitum access to food and water and a 12-hour light-dark cycle.

### Cisterna magna cannulation

The animal was anesthetized with 3% isoflurane mixed with 100% O_2_ in an induction chamber and transferred to a stereotaxic frame with the head secured by ear bars and a tooth bar. Anesthesia was maintained with 1.5% isoflurane in 100% O_2_ delivered via a nose cone. The body temperature was maintained at ∼37°C with a heating tape attached to the surface of the stereotaxis frame. A midline dorsal neck incision was made to expose the dura mater. A small durotomy was made using a 30-gauge needle to expose the cisterna magna. A polyethylene micro-catheter (0.13 mm ID × 0.25 mm OD, Scientific Commodities, Lake Havasu City, AZ, US) was inserted into the intrathecal space and secured with cyanoacrylate glue. Tracer delivery via cisterna magna was first confirmed with bright-field microscopy of Evans blue and cryoimaging of CF594 hydrazide and FITC-dextran (Fig. S3).

### MRI protocol

MRI studies were performed on a horizontal bore 9.4T preclinical scanner (Bruker Biospin, Billerica, MA, US) using a 35-mm volume coil. Mice were randomly assigned to three groups with intracisternal infusion of 1) 12.5 mM Gd-DTPA (n=7; MW=661.8 Da; Mallinckrodt, St Louis, MO, US); 2) 4.17 mM GadoSpin-P (n=6; MW=200 kDa; Miltenyi Biotec, Bergisch Gladbach, Germany); and 3) 90% enriched H_2_^17^O (n=7; MW=19 Da; NUKEM Isotopes, Alzenau, Germany). After the cannulation, the mouse was transferred to an MRI-compatible cradle and placed in a prone position. Anesthesia was maintained with 1 to 1.5% isoflurane delivered via a nose cone. Respiration rate and body temperature were monitored during the MRI scan. The respiration rate was maintained within 90 to 110 breaths per minute (bpm) by adjusting the anesthesia level. The body temperature was maintained at ∼37°C by blowing warm air into the scanner through a feedback control system (SA Instruments, Stony Brook, NY, US). After the initial setup, an anatomic scan was acquired at baseline using a 3D spin-echo sequence with the following parameters: TR/TE, 50/8.1 ms; FOV, 20×16×14 mm^3^; matrix size, 100×80×70; NAV, 2. Total scan time was ∼9 min. Subsequently, DCE-MRI acquisition was started, and 10 μL of contrast agent was infused at a rate of 1 μL/min (10 min total infusion time).

The transport of Gd-DTPA and GadoSpin was tracked dynamically using a T_1_-weighted 3D FLASH sequence with the following parameters: TR/TE, 50/2.5 ms; flip angle, 15°; FOV, 20×16×14 mm^3^; matrix size, 100×80×70; NAV, 1, leading to an isotropic resolution of 200 μm and a temporal resolution of 5 min. A single baseline scan and 25 dynamic scans were acquired before, during, and after contrast agent infusion for 130 min.

H_2_^17^O transport was imaged with a T_2_-weighted multi-slice RARE sequence. A total of 28 slices were acquired with TR, 2500 ms; FOV, 20×16 mm^2^; matrix size, 100×80; slice thickness, 0.5 mm. 8 echoes were acquired in each TR, leading to an effective TE of 31 ms. Acquisition time for a single-average dataset was 25 s. A total of 312 images were acquired continuously for 130 min with the first 12 images acquired as baseline.

To evaluate the significance of H_2_^17^O exchange across the blood-brain barrier (BBB), blood was collected from a subset of mice infused with H_2_^17^O at the end of DCE-MRI scan (n=4). Additional experiments were performed on the bench to collect blood from mice 10 min after the completion of H_2_^17^O infusion (i.e., 20 min from the start of H_2_^17^O infusion, n=3), as well as from mice without H_2_^17^O infusion (n=3). The blood samples were centrifuged at 2400 RPM for 20 min and the plasma was extracted. The T_2_ values of the plasma samples were measured using a single-slice Car-Purcell-Meiboom-Gill (CPMG) sequence with the following acquisition parameters: TR, 10 s; FOV, 20×20 mm^2^; matrix size, 128×128; slice thickness, 1 mm. A total of 64 echoes were acquired with an evenly spaced echo time of 8 ms.

### MRI image analysis

All images and data analyses were performed using either in-house developed or open-source software in MATLAB (MathWorks, Natick, MA, US) or Python (Python Software Foundation, v.3.0). Single-average T_1_-weighted images were reconstructed to delineate the kinetics and distribution of Gd-DTPA and GadoSpin transport. For the analysis of H_2_^17^O transport, T_2_-weighted images were reconstructed using 6 averages, resulting in a temporal resolution of 2.5 min. Motion correction was performed by registering the dynamic images to the anatomic image acquired at baseline via affine transformation using the open-source toolkit, Advanced Normalization Tools^35–37^.

Following motion correction, image segmentation was performed by co-registering images to an MRI mouse brain atlas^38^. Specifically, a representative animal was selected from each contrast agent group, and the anatomic images of the representative animal were registered to the atlas through affine and deformable transformation. Subsequently, images from the remaining animals in each contrast agent group were registered to that of the representative animal, followed by the generation of averaged dynamic images and tMIPs.

A total of 20 ROIs were generated from the co-registered brain atlas, covering the intracisternal infusion and transport pathways, the brain parenchyma, and the lateral ventricles. Mean signal intensity in each ROI was calculated, followed by the subtraction of the baseline signal for the entire dynamic series. Subsequently, the maximal signal from a small region surrounding the infusion catheter in the cisterna magna was used as the “input function” to normalize the time course of signal changes in each ROI.

### Clustering analysis

A correlation-matrix-based hierarchical clustering method was used to analyze the time courses of Gd-DTPA and H_2_^17^O transport among different ROIs^39^. Specifically, time-lagged cross-correlation analysis was performed to determine the similarity of the kinetics of contrast agent transport between two ROIs, as well as the lag time corresponding to the maximal cross-correlation coefficient (mCC)^40^. Subsequently, the “dissimilarity”, defined as 1-mCC, was used to quantify the distance between two ROIs. A dendrogram representing the hierarchical structure of the mCC matrix with complete linkage was generated using the open-source library SciPy^41^. Bootstrap analysis using the Pvclust package was performed to assess the uncertainty in clustering analysis^42^. A bootstrap replication of 1000 was used to calculate the approximately unbiased (AU) p-values and the bootstrap probability (BP) values. Visualizations of ROIs in each cluster were created using the Allen mouse brain atlas and brainrender^43,44^.

## Declarations

## Acknowledgments

The author(s) would like to thank Drs. Junqing Zhu and Chunying Wu for their assistance in establishing the intracisternal infusion protocol in mice.

## Authors’ contributions

YZ, experimental design, technical development and implementation, data collection and analysis, manuscript development and editing; GW, data analysis; CK, data collection and analysis; YG, technical development and implementation; HG, technical development; JZ, data analysis; XZ, data analysis; YW, technical development; DW, technical development; CF, experimental design, technical development; XY, experimental design, technical development, data collection and analysis, manuscript development and editing.

## Availability of data

Experimental data, images, and code from this study are available upon request to the corresponding author.

