## Supplemental figures for "Transport Pathways and Kinetics of Cerebrospinal Fluid Tracers in Mouse Brain Observed by Dynamic Contrast-Enhanced MRI"

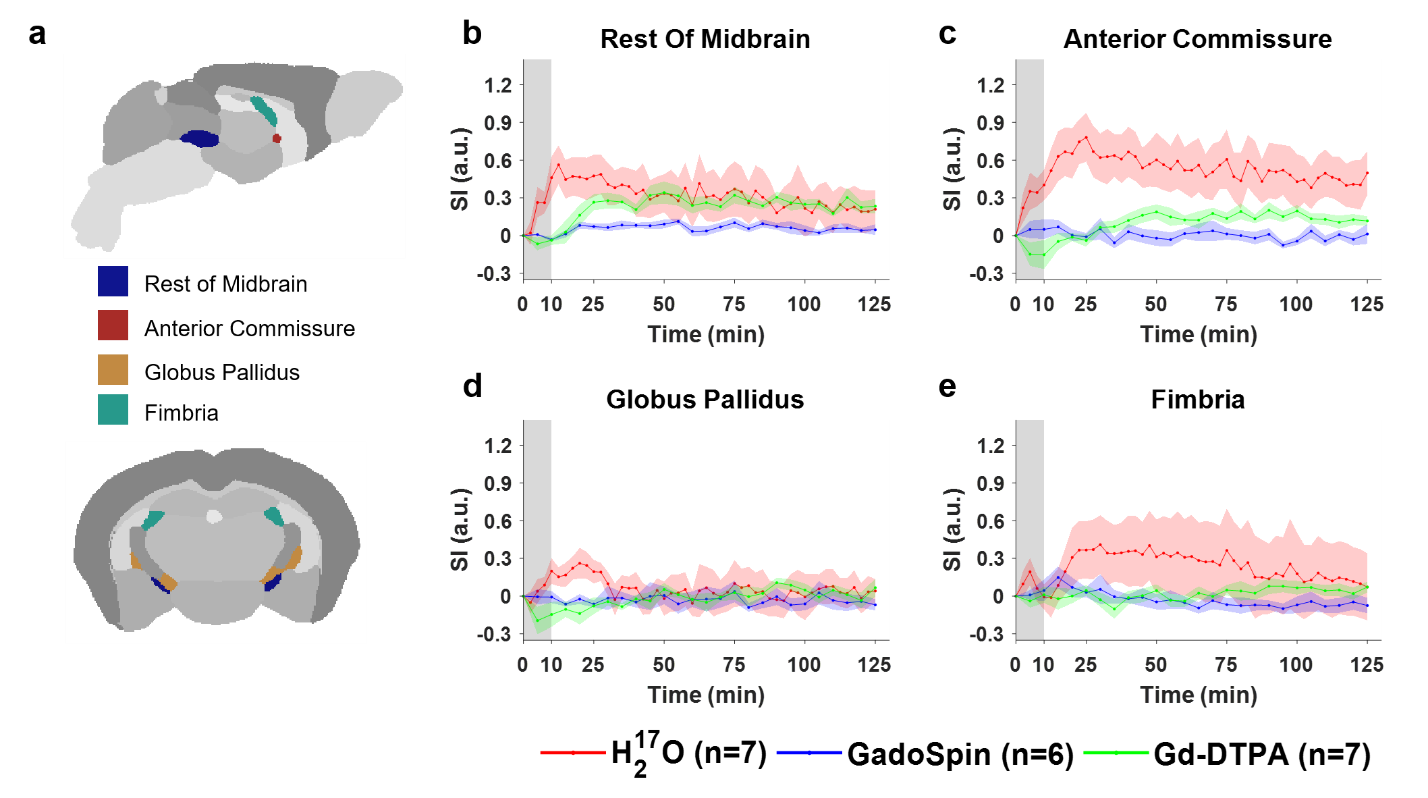

**Fig. S1** Contrast agent transport in the rest of midbrain, anterior commissure, globus pallidus, and fimbria. **a:** Segmentation of selected ROIs. **b-e:** Time courses of signal changes in the selected ROIs. Gray bands indicate the time period of contrast agent infusion. Red, blue, and green lines represent the mean time courses of signal changes induced by H_2_^17^O, GadoSpin, and Gd-DTPA, respectively. Shaded areas represent standard errors.

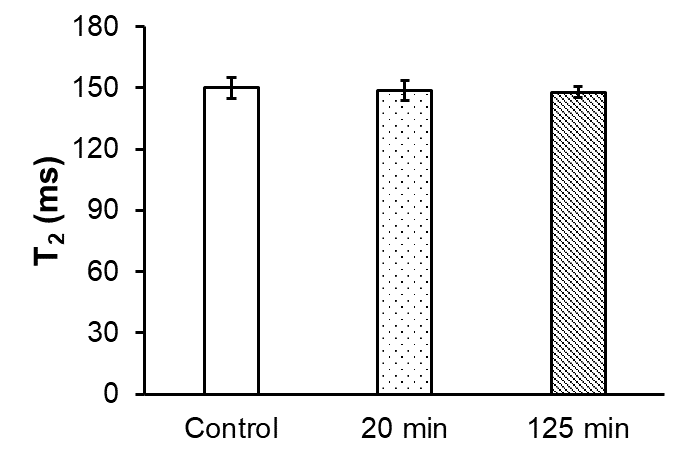

**Fig. S2** T_2_ of plasma from mice without H_2_^17^O infusion (control) and at 20 and 125 min after H_2_^17^O infusion.

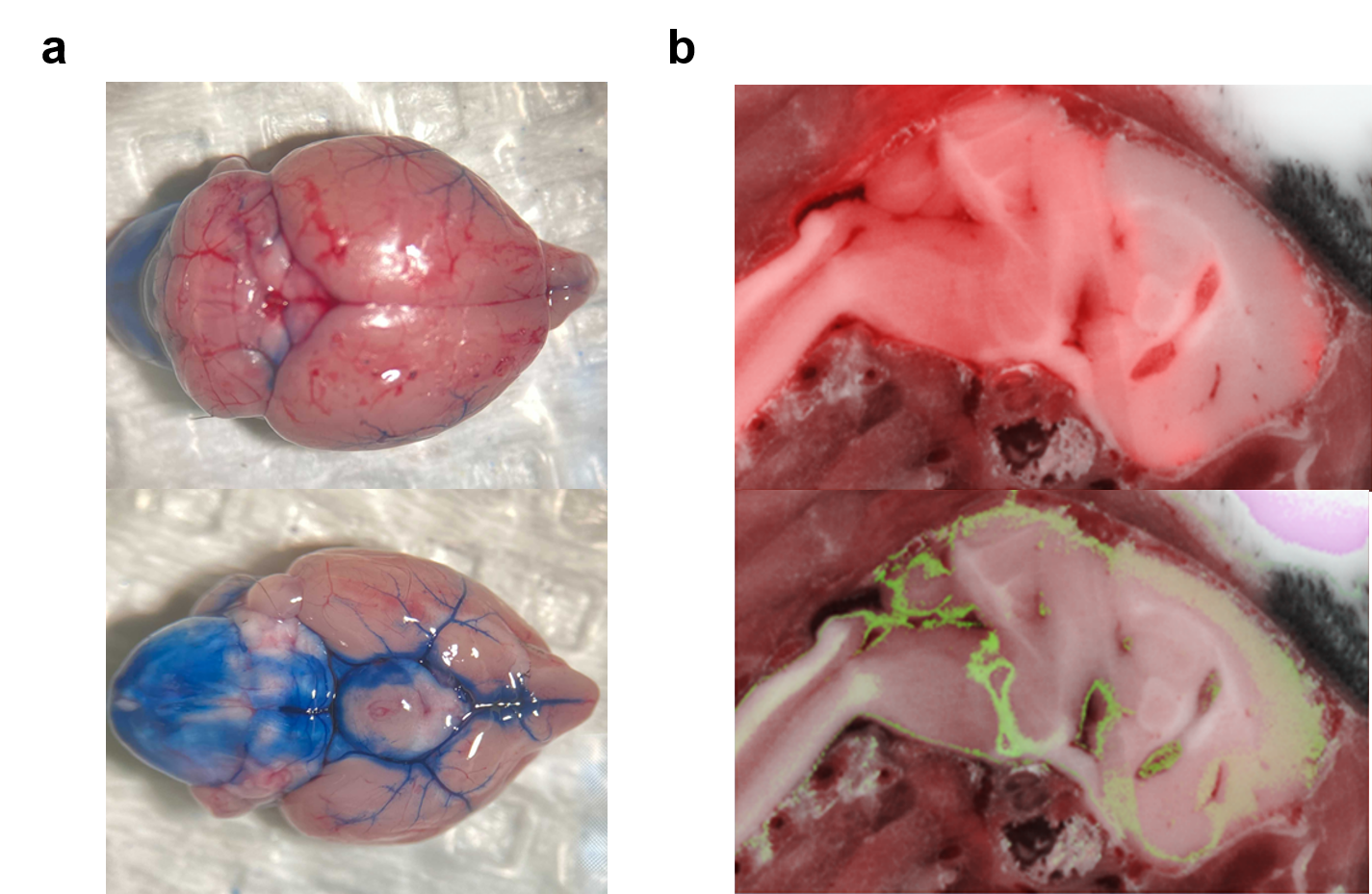

**Fig. S3** Validation of intracisternal tracer delivery. **a:** Bright-field microscopy of Evans blue staining. Top: dorsal view; bottom: ventral view. **b:** Cryoimaging of co-injected CF594 hydrazide (MW=740 Da, red, top) and FITC-dextran (MW=2,000 kDa, green, bottom). CF594 hydrazide showed penetration into the parenchyma while Evans blue and FITC-dextran were confined to the subarachnoid and perivascular spaces.
